# Disordered yet functional atrial t-tubules on recovery from heart failure

**DOI:** 10.1101/2021.10.28.466162

**Authors:** Jessica L. Caldwell, Jessica D. Clarke, Charlotte E.R. Smith, Christian Pinali, Callum J. Quinn, Charles M. Pearman, Aiste Adomaviciene, Emma J. Radcliffe, Amy E. Watkins, Margaux A Horn, Elizabeth F. Bode, Mark Eisner, David A. Eisner, Andrew W. Trafford, Katharine M. Dibb

## Abstract

Transverse (t)-tubules drive the rapid and synchronous Ca^2+^ rise in cardiac myocytes. The virtual complete loss of atrial t-tubules in heart failure (HF) decreases Ca^2+^ release. It is unknown if or how atrial t-tubules can be restored and if restored t-tubules are functional.

Sheep were tachypaced to induce HF and recovered when pacing was stopped. Serial block face Scanning Electron Microscopy and confocal imaging were used to understand t-tubule ultrastructure and function. Candidate proteins involved in atrial t-tubule recovery were identified by western blot and causality determined using expression studies.

Sheep atrial t-tubules reappeared following recovery from HF. Despite being disordered (branched, longer and longitudinally arranged) recovered t-tubules triggered Ca^2+^ release and were associated with restoration of systolic Ca^2+^. Telethonin and myotubularin abundance correlated with t-tubule density and altered the density and structure of BIN1-driven tubules in neonatal myocytes. Myotubularin had a greater effect, increasing tubule length and branching, replicating that seen in the recovery atria.

Recovery from HF restores atrial t-tubules and systolic Ca^2+^ and myotubularin facilitates this process. Atrial t-tubule restoration could present a new and viable therapeutic strategy.

**Brief Summary:** The loss of atrial transverse (t)-tubules and the associated dysfunction in heart failure is reversible and the protein myotubularin plays an important role.

## Introduction

Transverse (t)-tubules are invaginations of the surface membrane of ventricular myocytes containing ion channels and regulators of excitation contraction coupling which synchronize the systolic rise of [Ca^2+^]_i_ (1). Ventricular t-tubule loss occurs in various cardiovascular diseases reducing the amplitude and synchronicity of the systolic Ca^2+^ transient (1, 2). The extent of t-tubule remodeling correlates with left ventricular dysfunction (3). Large mammals, including humans, have extensive atrial t-tubule networks which are reduced in heart failure (HF) and atrial fibrillation (AF) (4–7). However, in contrast to the ~20% reduction in ventricular t-tubules, HF causes near-complete loss of atrial t-tubules with profound effects on systolic Ca^2+^ (4).

T-tubules can be at least partially restored in the failing left ventricle by mechanical unloading (8), SERCA2a gene therapy (9), resynchronization therapy (10), treadmill exercise (11) β-blockers (12) and PDE5 inhibition (13). Some work also shows restoration of systolic Ca^2+^(8–10, 13, 14). However, since only ~20% ventricular t-tubules are lost (1, 7, 9), newly formed t-tubules cannot be distinguished from pre-existing t-tubules. Therefore, whether new t-tubules are functional, and whether the restoration of systolic Ca^2+^ is due to new t-tubules rather than enhanced Ca^2+^ release of pre-existing t-tubules remains unknown. This is essential to understand if t-tubule restoration has the potential to restore contractile function in disease. Uniquely, the near-complete loss of atrial t-tubules in HF (4, 7) allows us to determine if restoration is possible and if new atrial t-tubules can trigger Ca^2+^ release necessary for restoration of systolic Ca^2+^.

The process by which t-tubules are formed is largely unknown. A number of proteins have been implicated in the ventricle including junctophilin 2 (JPH2) (15), amphiphysin II (Amp-II, BIN-1) (7, 16) and telethonin (Tcap) (17) but any role in the atria remains to be determined. For t-tubule restoration to be therapeutically viable an understanding of this process is essential. Given these considerations, the main aims of this study were to establish; i) if new atrial t-tubules can form once they have been lost in HF, ii) if new atrial t-tubules can trigger Ca^2+^ release and, iii) which proteins drive formation of new atrial t-tubules.

## Results and Discussion

### Atrial t-tubules are restored following recovery from HF

As shown previously, HF is associated with ventricular dilatation and reduced contractility (4, 7, 13, 18)(Supplementary Table 1) accompanied by near-complete atrial t-tubule loss in HF (4) (Figure 1A&B). We now report that cessation of tachypacing reversed subjective signs of HF and partially restored ventricular contractility (Supplementary Table 1). Atrial cellular hypertrophy was reversed (Supplementary Table 1) and atrial t-tubule density fully recovered (Figures 1A&B) following termination of tachypacing, demonstrating t-tubule recovery post HF is possible. Therefore, our model provides an unparalleled opportunity to understand the mechanisms and functional consequences of restoring atrial t-tubules.

**Figure 1.**
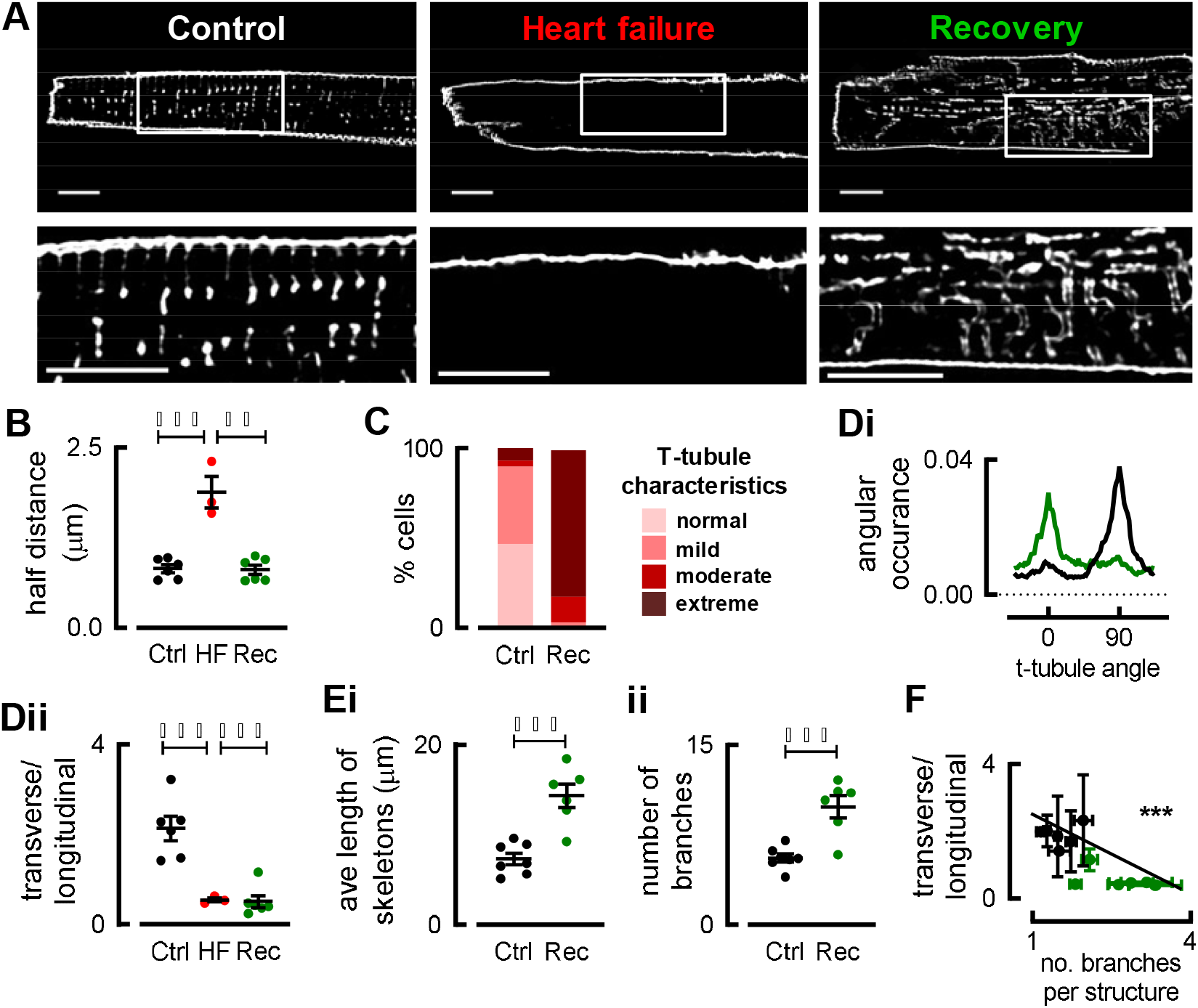
Sheep atrial t-tubules are restored following recovery from heart failure. (A) Di-4-ANEPPS staining of sheep a al myocytes showing t-tubule lo HF and restoration in recovery. (B) Mean half dist to nearest t-tubule and surface m m ane. (C) Categorization of t-tubule disorder. (D) T cal control (black) and recovery (green) t-tubule orientation (i); transverse tubules (90°) and longitudinal (0°) and (ii) mean transverse:longitudinal tubule ratio. (E) Mean data calculated from skeletonized images for t-tubule length (i) and branching (ii). (F) Relationship between branching and t-tubule angle; symbols denote animals (R^2^=0.614). n=24-61 cells, N=3-6 animals.**p<0.01,***p<0.001; linear mixed modelling. Scale bars=10μm.

### Recovered atrial t-tubules are disordered

Despite restoration of atrial t-tubule density, t-tubule organization is dramatically altered in atrial cells following recovery from HF (henceforth termed recovery cells; Figure 1A bottom right & 2B). Although HF causes t-tubule disorder in the ventricle (19–21), such disorder has not been reported in the atria or following reverse ventricular remodeling (8–11, 14). Quantification of t-tubule disorganization revealed that extreme disorder (supplemental methods) increased from 2% in control to 79% following recovery (blinded observations; Figure 1C). To characterize the underlying structural changes t-tubule images were skeletonized (7) and their orientation determined. In control sheep (similar to our work on human atrial tissue (5), although in contrast to smaller mammals (22, 23)) atrial t-tubules were mostly transversely orientated with few longitudinal components. However, substantial longitudinal components did appear following recovery from HF (Figures 1Di&ii), contributing to the t-tubule disorder. In addition, t-tubules in recovery cells were longer and more branched (Figure 1Ei&ii) and animals with the most t-tubule branching also contained more longitudinally-orientated tubules (Figure 1F;p<0.001).

**Figure 2.**
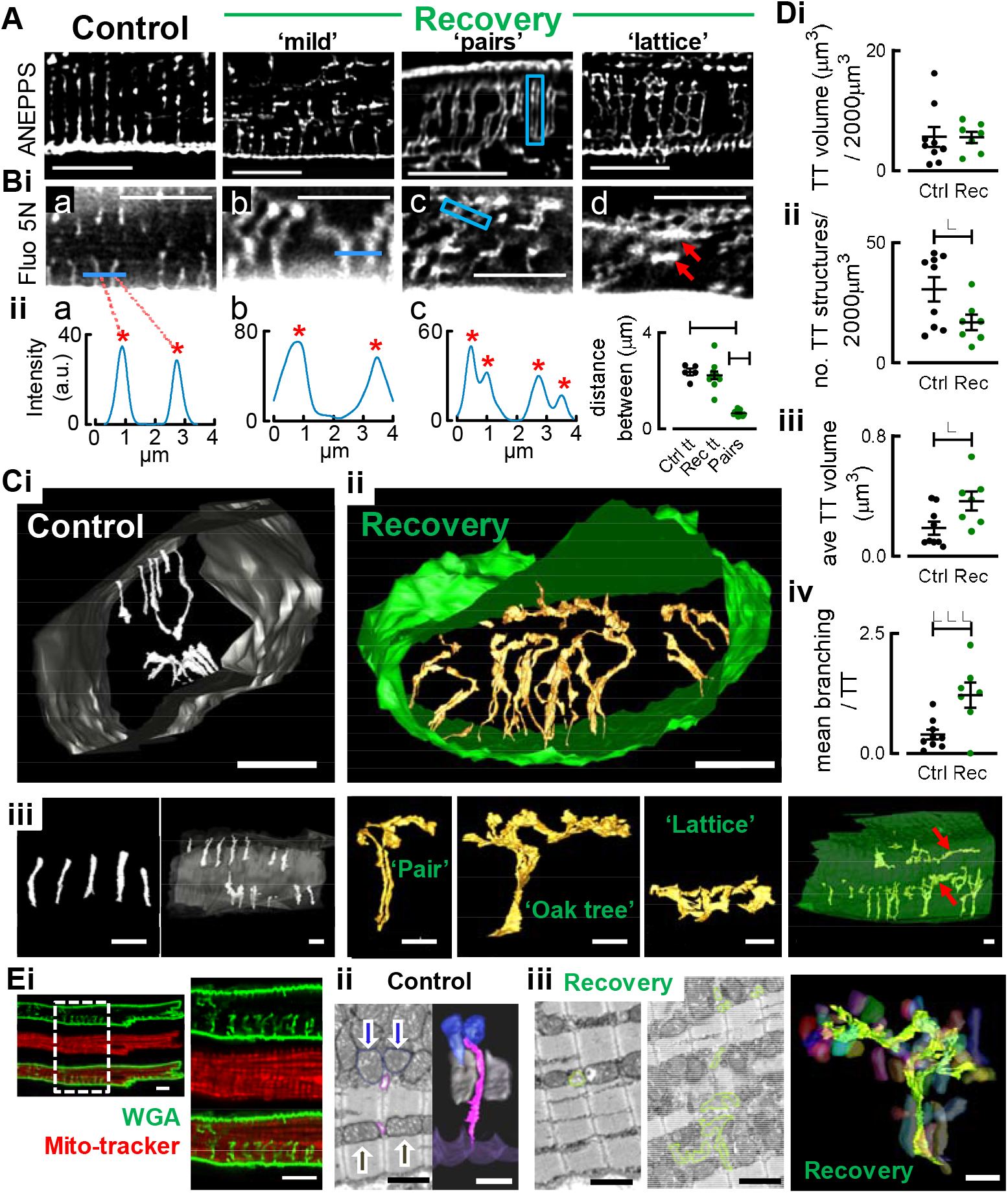
The diversity of atrial t-tubule remodeling. (A) Di-4-ANEPPS staining of control and recovered atrial myocytes. (Bi) Bath application of Fluo 5N showing t-tubules open to the cell exterior and can occur in pairs (blue box A&Bic). Tubule pairs were confirmed by intensity plots (Bii shows examples which correspond to intensity along blue lines/boxes in Bi and red asterisks denote peaks corresponding to t-tubules). Mean spacing of t-tubules and individual t-tubules forming pairs is shown in Bii; 15-42 tubules, n=5-11,***p<0.001;ANOVA. Scale bars=10μm. (C) Representative sbfSEM 3D reconstructions of control (i) and recovery (ii) atrial myocytes, scale bars=5μm. (iii)Common t-tubule morphologies and longitudinal cell views for control (left and recovery (right);scale bars=2μm. (D) Mean data calculated from sbfSEM for t-tubule volume and number per cell volume (i&ii), average t-tubule volume (iii) and average branching per t-tubule (iv). (Ei) Recovered atrial myocyte imaged confocally showing disordered t-tubules (green) weaving between mitochondria (red;scale=10μm). (Eii) Example 2D-section and model of a control atrial t-tubule (pink) extending along the z-line between a single band of mitochondria (grey arrows/mitochondria) but not penetrating the thicker mitochondrial bed (bue arrows/mitochondria; scale=2μm). (Eiii) Recovered atrial t-tubule (lime) which extends normally where mitochondria occur in narrow bands (left) but penetrates thick mitochondrial beds forming irregular branching (middle/right).*p<0.05,***p<0.001; students t-test; n=18, N=3 animals/group.

Recovered atrial t-tubule disorder varied between cells and animals ranging from ‘mild’ to extreme including tubule ‘pairs’ (blue box in Figure 2A) and ‘lattice’ like structures (Figure 2A). Experiments using a cell impermeant fluorescent Ca^2+^ indicator fluo-5N in the extracellular solution (Figure 2B) showed that it entered disorganized tubules indicating they open to the cell exterior and, being continuous with the surface membrane, are likely functionally important (Figure 2B). Figure 2Bid (red arrows) shows large dilated longitudinally-orientated tubules which, being continuous with the surface membrane, are also likely functional. Fluo-5N did not fill the space between parallel transverse tubule pairs (blue box Figure 2Bic) suggesting they are separate tubules in close proximity as opposed to a poorly resolved larger structure. Normal t-tubules were on average 2.22±0.16 μm apart (distance between red asterisks; Figure 2Biia,b&d). When t-tubules occurred in pairs, each ‘pair’ was similarly spaced however individual tubules forming pairs were only 0.65±0.03 μm apart (Figure 2Biic-d) indicating non z-line localization. Therefore, recovered t-tubules are often highly disordered, longitudinally-orientated, and can be very wide, paired and lattice like in appearance. Since confocal microscopy lacks the resolution to understand the ultra-structure of these tubules we used serial block-face scanning electron microscopy (sbfSEM).

### Recovered atrial t-tubules are large and exhibit extreme structural disorder associated with disrupted mitochondrial positioning

Sheep atrial t-tubules occupy ~0.27% cell volume, project radially from the sarcolemma towards the cell interior (as in human atrial myocytes (5)), occur primarily at z-lines and do not contact neighboring tubules. In contrast to the relatively uniform appearance of ventricular t-tubules (24), control atrial t-tubules are morphologically heterogeneous (Figure 2Ci, Supplemental Figure 1). Some common morphologies (‘column’ and ‘angled’) are shared with ventricular t-tubules, but others (‘stump’ and ‘club’) appear unique to the atria (Figure 2C; t-tubule morphologies are categorized in Supplemental Figure 1) and could influence dyad formation.

Consistent with confocal data, recovered atrial t-tubules are highly disordered, often spanning several sarcomeres (see red arrows in Figure 2Ciii right and also by the oak-tree and lattice); normal t-tubule morphologies are replaced by disordered structures e.g. random, oak-tree, longitudinal, lattice and pairs which together account for 61% of the total t-tubule volume in recovery (Supplemental Figure 1A&B). Compared to control, there are fewer t-tubules in recovery but their volumes were larger, in line with increased tubule branching, resulting in complete restoration of t-tubule density (Figure 2D). This is the first report of such atrial t-tubule remodeling and is more extreme than that seen in the ventricle e.g. (7, 24). We next sought to determine if t-tubule remodeling is associated with structural remodeling of other cellular components.

Atrial mitochondria appear disrupted following recovery from HF and confocal data suggests that disordered t-tubules weave between mitochondria (Figure 2Ei). Control atrial t-tubules extend between rows of mitochondria which are 1 mitochondrion deep (grey arrows/mitochondria Figure 2Eii) but do not penetrate wider mitochondrial beds (blue arrows/mitochondria). Following recovery from HF sbfSEM shows enlarged mitochondrial beds; here mitochondria are irregularly arranged creating large gaps occupied by debris (Figure 2Eiii middle). Recovered t-tubules can extend normally along the z-line to these regions (with single rows of mitochondria; Figure 2Eiii LHS) but recovered t-tubules enter wide, disordered mitochondrial beds and occupy the space, becoming highly branched and dilated and allowing the formation of ‘oak-tree’ structures (Figure 2Eiii middle;lime tracing). Following recovery, mitochondria are also present on the z-line where they seem to block normal t-tubule extension causing branching (Supplemental Figure 2). Our data suggests an association between t-tubule and mitochondria disruption whereby disordered mitochondrial beds allow erratic t-tubule extension and misplaced mitochondria may divert the normal t-tubule extension along the z-line. Since triggered Ca^2+^ release relies on a close association between t-tubules and the SR, we next sought to determine if disordered atrial t-tubules can trigger Ca^2+^ release in the cell interior.

### Despite their structural disorder recovered atria t-tubules still trigger Ca^2+^ release

Importantly, here we show HF-associated dysfunction can be reversed as recovery from HF completely restored the atrial Ca^2+^ transient amplitude in voltage clamped cells (Figure 3Ai-ii). To establish if triggered Ca^2+^ release from newly formed, recovered t-tubules plays a role, Ca^2+^ was recorded using high-speed XY confocal imaging (Figure 3B). The early rise of [Ca^2+^]_i_ was overlaid with the t-tubule/surface membrane staining (WGA) from the same confocal plane (Figure 3Ci-iii). The rise of systolic [Ca^2+^]_i_ occurs first along the t-tubules in control whilst in HF cells, where t-tubules are absent, it is restricted to the cell surface (Figure 3Ciii). Following recovery, [Ca^2+^]_i_ also rises first along disordered t-tubules *before* propagating to the rest of the cell (Figure 3Ciii) showing new t-tubules trigger Ca^2+^ release.

**Figure 3.**
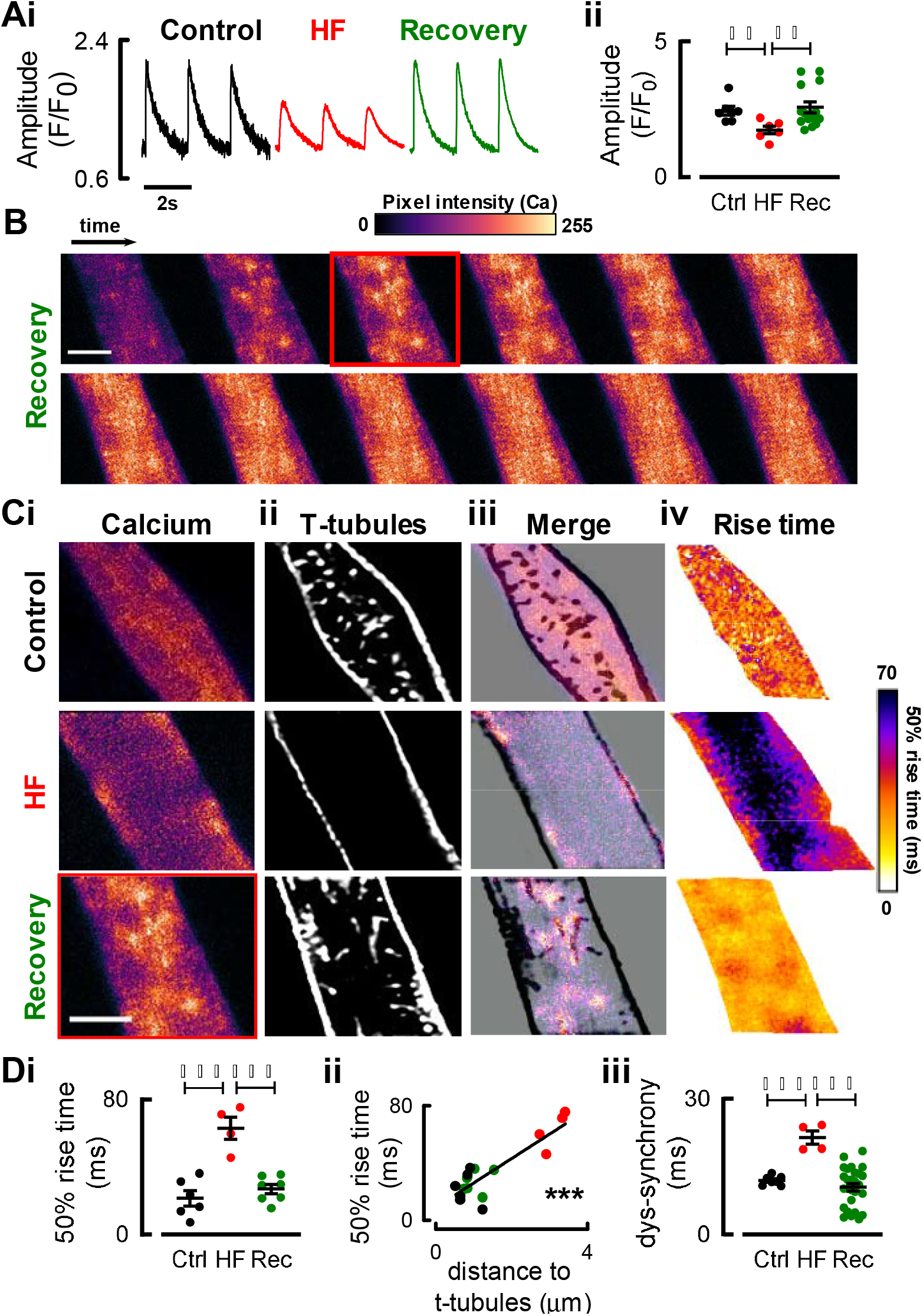
Recovered t-tubules trigger Ca^2+^ release and are associated with restored ystolic Ca^2+^. (A) Representative Ca^2+^ transi s and mean systolic Ca^2+^ transient mplitude. (B) Representative time series from recovery atrial cell showing early calc um release (fluorescence). (C) Representati atrial myocytes showing; triggered alcium release (i), membrane staining (ii), merg of i and ii (iii) and Ca^2+^ rise time (iv). ummary data for 50% Ca^2+^ rise time (Di), corre tion between rise time and distance t t-tubules (R^2^=0.8457;Dii) and dys-synchrony Ca^2+^ release (Diii). n=4-14 cells, N= animals for recovery and N=1-5 animals/grou (control and HF to confirm our previous work (4).***p<0.001;ANOVA. Scale bars=10μm.

The loss of atrial t-tubules in HF slows the rise of Ca^2+^ (Figures 3Civ & Di). However, despite t-tubule disorder, the rate of rise of Ca^2+^ returned to control values in the recovery cells due to the restoration of atrial t-tubules (Figures 3Civ & Di-ii). Therefore, recovered t-tubules are as effective as control t-tubules in triggering calcium release. In the ventricle, dys-synchronous Ca^2+^ release arising from t-tubule loss (25) could promote arrhythmias (26) and therefore recovery from dys-synchrony is beneficial. In HF, atrial t-tubule loss also increased dys-synchrony but this was fully corrected upon recovery (Figure 3Diii) presumably since new t-tubules trigger Ca^2+^ release in the cell interior. Thus, despite their disorder, recovered t-tubules restore triggered Ca^2+^ release suggesting atrial t-tubule recovery is a future novel and viable therapeutic approach.

### Increased myotubularin in the recovery atria may underlie increases in t-tubule density, length and branching

It is important to understand how new t-tubules are built. Since recovered atrial t-tubules are almost entirely new structures, to better understand their formation we investigated how the abundance of proteins implicated in ventricular t-tubule formation changed with atrial t-tubule loss and recovery (Figure 4A). BIN1 tubulates ventricular neonatal cells which lack t-tubules (13) (Figure 4B), and its expression correlates with t-tubule density (7, 9). However, in the atria BIN1 decreased modestly in HF but upon recovery was not different to control or HF suggesting other proteins are also involved (Figure 4A). Our data also suggest changes in atrial t-tubule density do not involve junctophilin 2 (JPH2). However, telethonin (Tcap), a load-dependent regulator of ventricular t-tubules in some but not all models (9, 13, 17), correlated with atrial t-tubule abundance (Figure 4A). Additionally, the BIN1 binding partner and regulator (in skeletal muscle) myotubularin (MTM1 (27)), which to the best of our knowledge has not been shown to regulate cardiac t-tubules, also correlates with atrial t-tubule density (all P < 0.05). We next investigated if increased expression of Tcap and MTM1, as occurs in recovery from HF, can *directly* alter t-tubule density or structure.

**Figure 4.**
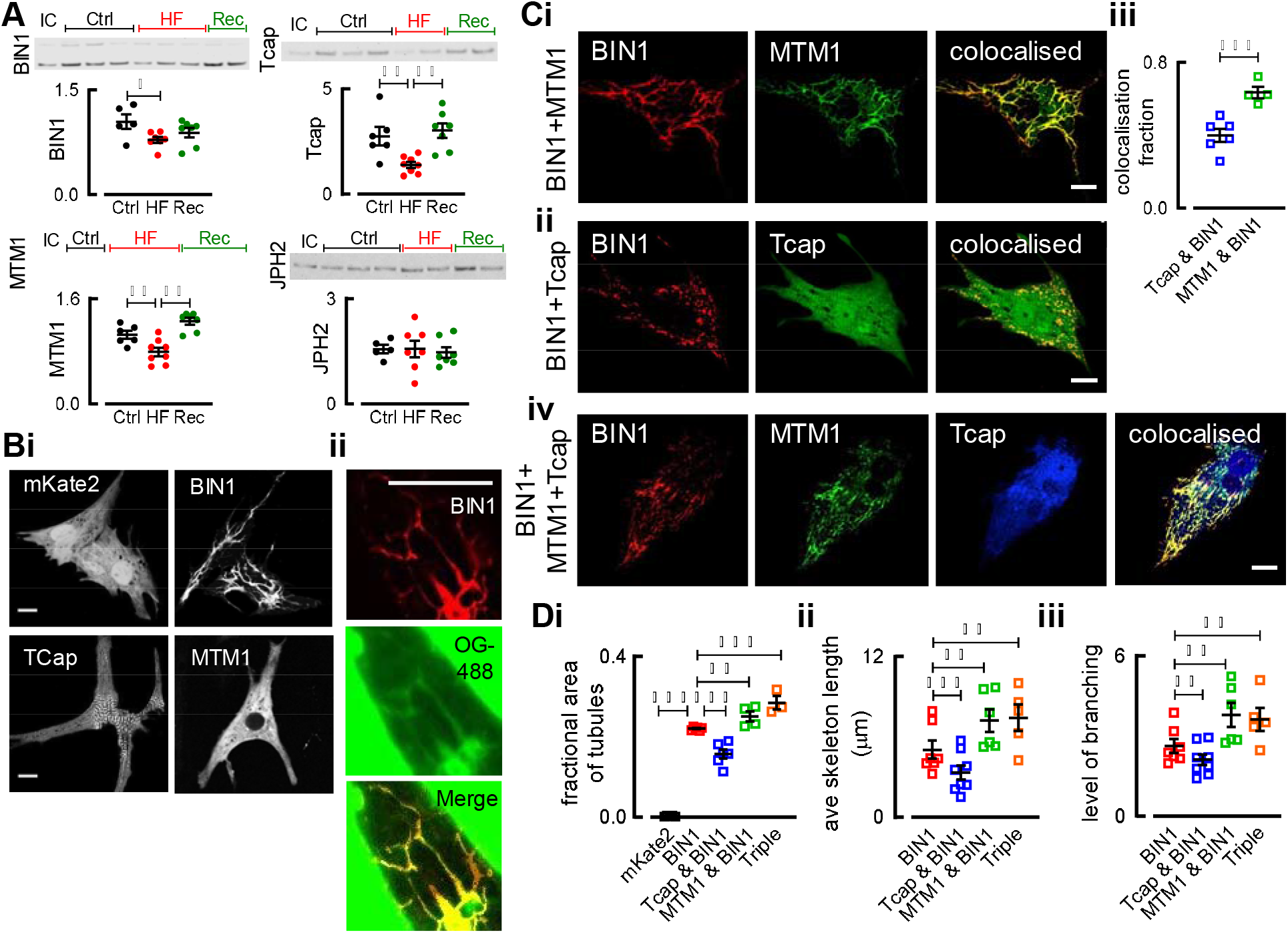
MTM1 and Tcap influence t-tubule density an structure and MTM1 likely facilitates atrial t-tubule restoration. (A) Represen tive western blots and mean data for BIN1, Tcap, MTM1 and JPH 2 in control, HF a d recovery sheep atrial tissue; N=7,*p<0.05,**p<0.01 (ANOVA;nested statistics). (Bi) Representative NRVMs transfected with mKate2 (empty vector), BIN1, MTM1 or Tca xpression vectors. (Bii) Oregon Green (green) staining of BIN1 transfected cell (red) and merge. (C) Representative NRVMs transfected with BIN1 (Red), MTM1 (Green; Ci) or Tcap (Green; Cii) or triple transfection of BIN1 (Red), Tcap (Blue) and MTM1 (Green; Civ). Colocalisation (yellow) summarized in Ciii. (D) Mean data summarizing; fractional area of cells occupied by tubules (i); average skeleton length (ii); average branching of each structure (iii). N=5-8 isolations, n=45-98 cells.*p<0.05,**p<0.01,***p<0.001 (ANOVA;linear mixed modeling). Scale bars=10μm.

Neonatal rat ventricular myocytes (NRVMs) lack t-tubules (Figure 4B). While the control vector, MTM1 or Tcap alone failed to produce tubules, expression of BIN1 did (Figure 4Bi&Di). BIN1 driven tubules were connected to the extracellular environment as shown by extracellular dye penetration (Oregon Green) which colocalized with BIN1 (Figure 4Bii).

We co-expressed BIN1 with either Tcap, MTM1 or both to elucidate the role of MTM1 and Tcap in modulating BIN1 tubulation (Figure 4Ci, ii&iv). MTM1 colocalized with BIN1 at tubules, however there was little colocalization between BIN1 and Tcap (Figure 4Ciii). Tubule structures generated by transfection of BIN1 with MTM1, Tcap or both are shown in red (Figure 4C LHS). Tubule density was decreased by Tcap but increased by MTM1 (Figure 4Di). Atrial t-tubule recovery is associated with increases in both Tcap and MTM1; the triple transfection (BIN1+MTM1+Tcap; Figure 4Civ) increased t-tubule density compared to BIN1 alone suggesting the MTM1 effect is dominant (over Tcap) and may underlie the increase in atrial t-tubule density upon recovery from HF.

Since the recovery atria had longer, branched tubules, we next investigated if either protein could alter the structure of BIN1-driven tubules. Both the average skeleton length (equivalent to the length of a single tubule structure) and level of branching were decreased with Tcap but increased with MTM1 (Figure 4Dii&iii). Again, the MTM1 effect was dominant in the triple transfection for both parameters suggesting MTM1 plays a role in the increased atrial t-tubule length and branching following recovery from HF. Therefore, taken together, we have shown for the first time that MTM1 can modulate cardiac t-tubule density and structure. We suggest increased MTM1 in the recovered atria is important for the increase in t-tubule density, length and branching and may therefore represent a future therapeutic target in the atria.

### Atrial RyRs move away from the z-line in HF but z-line localization is restored following recovery from HF

BIN1 and MTM1 have been suggested to alter dyadic structure and RyR localization in cardiac and skeletal muscle respectively (28, 29). Our observation that disordered, recovered t-tubules can still trigger Ca^2+^ release suggests functional dyads containing L-type Ca^2+^ channels (LTCC) and RyRs are formed. This is supported by Na^+^-Ca^2+^ exchanger (NCX) localization at both the normal and disordered t-tubules (Supplemental Figure 3A).

RyRs occur on the z-line in control and recovery, colocalizing with t-tubules (Supplemental Figure 3Bi-iii). Surface RyRs were sparse in control and recovery but increased in HF (Supplemental figure 3Biv). Since RyR density does not increase in HF (18), this suggests RyRs shift from the z-line to the surface, allowing triggered Ca^2+^ release at the surface. Compared to control and recovery, where RyR staining appears continuous along the z-line, in HF RyR staining is fragmented and occurs between z-lines (Supplemental Figure 3C). We speculate RyR fragmentation contributes to the dys-synchronous Ca^2+^ release and RyRs between z-lines may promote Ca^2+^ wave propagation (30). Decreased MTM1 and BIN1 may play a role in atrial RyR disorder in HF (28, 29).

### Conclusion

We show, for the first time that atrial t-tubules can be restored following their almost complete loss in HF. Although these new t-tubules are highly disordered, they can trigger Ca^2+^ release in the cell interior and completely recover the amplitude of the systolic Ca^2+^ transient and synchrony of Ca^2+^ release. Whilst Tcap and MTM1 influence BIN1 driven tubule density and structure, MTM1 is dominant and we suggest it facilitates restoration of t-tubule density by producing longer, more branched atrial t-tubules. Restoring atrial t-tubules by targeting MTM1 may be possible in the future in diseases such as atrial fibrillation and HF in which t-tubules are lost.

## Methods

For detailed Methods see the online supplement.

The study accords with The United Kingdom Animals (Scientific Procedures) Act, 1986, European Union Directive (EU/2010/63) and is reported in accordance with the ARRIVE guidelines. Institutional approval was obtained from The University of Manchester Animal Welfare and Ethical Review Board.

## Author Contributions

Authors contributed to the delivery of experiments/data analysis. In addition, KMD, DAE & AWT designed experiments and prepared the manuscript.

## Acknowledgements

This work was supported by research grants from The British Heart Foundation (FS/12/34/29565,PG12/89/29970,FS/14/4/30532,PG/18/24/33608). The authors acknowledge the technical support for high rate pacing protocols and devices provided by Medtronic (USA).

The authors thank the staff in the FBMH EM Core Facility for their assistance and the Wellcome Trust for equipment grant support to the EM Facility.

