## Supplemental Data for "Disordered yet functional atrial t-tubules on recovery from heart failure"

### Methods

All procedures involving animals accord to the United Kingdom (Scientific Procedures) Act of 1986 and the University of Manchester Ethical Review Board.

***Tachypacing induced heart failure in the sheep* –** 37 female Welsh Mountain sheep (, ~ 18 months of age) were randomly assigned to control (non-instrumented), heart failure (HF) or recovery groups. Under isoflurane anaesthesia (1–4% v/v in oxygen) 25 sheep were instrumented with a pacing lead (St Jude Medical or Medtronic), fixed transvenously at the right ventricle apex and attached to a subcutaneous cardiac pacemaker (Medtronic). Sheep were given post-operative analgesia (meloxicam 0.5 mg/kg) and antibiosis (enrofloxacin 2.5 mg/kg). Animals recovered post-operatively for one week before tachypacing was initiated. HF was induced by rapid ventricular pacing (210bpm; median 35 days, interquartile range 12.5 days) as described previously (1-3). At the point of HF pacing was terminated in a sub-set of sheep that were allowed to recover for 37.±1.2 days. Animals were monitored daily for onset of clinical signs of HF (lethargy, dyspnoea and weight loss) and cardiac function was assessed regularly in conscious animals by trans-thoracic echocardiography as described previously (2-5).

***Isolation of sheep atrial cardiac myocytes -*** Sheep were killed (heparin 10,000 units and pentobarbitone 200 mg kg−1 intravenously), the heart excised and the ventricles and atria were separated. The left circumflex coronary artery of the left atria was cannulated and left atrial myocytes isolated using a collagenase and protease digestion technique as described previously (1, 3).

***Isolation of neonatal rat ventricular myocytes, cell culture and transfection -*** Ten litters (total) of two day old Wistar rats (Charles River UK ltd) were killed by cervical dislocation and decapitation. Bodies were rinsed in 70% ethanol and hearts excised and placed in ice cold disassociation buffer (containing (mM): 116 NaCl; 5.6 glucose; 20 HEPES; 5.4 KCl; 0.83 MgSO_4_; 1 NaH_2_PO_4_; (pH 7.35)). Single ventricular myocytes were isolated using warmed dissociation buffer, containing 0.75mg/ml Collagenase A (Roche), and 1.3mg/ml Pancreatin (Sigma), spun at 120 rpm at 37°C for 7 minutes followed by titration. Serial digestions were performed until the ventricles were completely digested. Myocytes were re-suspended in pre-plating media (containing; 68% DMEM, 17% Medium 199, 5% FBS, 1% fungizone, 1% penicillin streptomycin (penstrep; 10,000 units Pen/10mg/ml Strep) (Gibco, Life Technologies) and 10% horse serum (Sigma), plated on tissue culture dishes and left to settle for 75 minutes to allow fibroblasts to attach. The supernatant, containing NRVMs, was then removed and plated onto tissue culture dishes (Ibidi, GmbH) at a density of 4x10^5^ cells/ml in maintenance media (68% DMEM, 17% M199, 5% FBS, 1% fungizone, 1% penicillin streptomycin (10,000 units/ml) (Gibco, Life Technologies), 10% horse serum and 100µM Bromodeoxyuridine (Sigma) and maintained in a 5% CO_2_ incubator at 37°C.

NRVMs were transiently transfected 2-4 days after isolation with either human BIN1 (variant 8), MTM1 or Tcap (Origene Inc) cloned into pCMV6 fluorescent (mKate2, mGFP or mBFP respectively) entry vectors or with pCMV6-mKate2 (Origene Inc) as a negative control. Cells were transfected with a 1:3 ratio of Plasmid DNA (6 µg) to transfection reagent (Fugene 6; Promega) in reduced serum media (OptiMEM; Life Technologies, UK) for 48 hours. NRVMs that had been successfully transfected, thus expressing the fluorescent pCMV6 tag, were imaged using a Nikon A1R^+^ confocal microscope (excitation, 405, 488 or 561 nm; emission, 425-475, 500-530 or 553-618 nm). Nikon Elements imaging software, Manders overlap function, was used to determine the colocalisation of BIN1 with Tcap and MTM1.

***T-tubule imaging and analysis –*** Atrial myocytes were stained with 4 µM di-4-ANEPPS (Molecular Probes). Additionally, 300µM Fluo 5N (Molecular Probes) staining was used to fill atrial t-tubules and 100µM Oregon Green (Molecular Probes) staining used to fill NRVM tubules from the bathing solution. T-tubule imaging was performed on either a Leica SP2 confocal microscope at an x-y resolution of 100 nm and vertical z stacks of 162 nm (excitation, 488 nm; emission >515 nm) or a Nikon A1R^+^ confocal microscope at 100 nm xyz pixel dimensions (excitation, 488 nm; emission 500-530 nm). Where necessary 2 mM EDTA was added to avoid cell movement during the z-stack.

Following image acquisition, confocal stacks were deconvolved, using either Huygens Professional (Scientific Volume Imaging, Netherlands) or NIS elements (Nikon) software. Deconvolution was achieved using the point spread function (PSF) of the microscope. The PSF was calculated ‘theoretically’ (NIS elements, Nikon) using known microscopic parameters or ‘measured’ (Huygens Professional, Scientific Volume Imaging) using 100 nm diameter polystyrene beads (Molecular Probes, ThermoFisher Scientific), as described previously (1, 5, 6). Average fluorescence intensity was then determined from central sections of each cell imaged and confocal stacks thresholded (Image J, NIH).

T-tubule density in sheep atrial cells was assessed in two ways: 1) the fractional area of the cell occupied by t-tubules, calculated from binary converted images, where tubules were represented as a fraction of cell area (Image J, NIH)(5); 2) the distance at which 50% of voxels within the cell are from cell membrane (t-tubule and surface sarcolemma), known as half distance, was calculated using routines written in IDL (Exelis, UK). Distance maps were then calculated whereby, colour intensity represented the distance of each voxel of the cell was from the nearest membrane (Image J, NIH), as described previously (1, 5, 6).

To assess t-tubule disorder in sheep atrial cells, t-tubules were firstly categorized visually (blinded observations) into normal, mild, moderate or extreme disorder. T-tubule disorder included longitudinal orientation, branching or tubule pairs i.e. deviation from the normal transverse structures located on the z-line. T-tubule orientation was determined using binary images which were “skeletonized” and the Fiji (Image J, NIH) plug-in “directionality” used to generate frequency plots of t-tubule angles (5, 7). To quantify t-tubule orientation, the sum of t-tubules corresponding to two cell directions (longitudinal versus transverse) was measured and the ratio between the two calculated (5). Skeletonised images were also used to obtain t-tubule branch information where the Fiji plug-in “analyse skeleton” calculated the average length and the number of elements a tubule structure was comprised of.

Tubule structure was assessed in NRVMs using the Fiji (Image J, NIH) plug-in “Ridge detection” to firstly define tubules (lines of less than 2um were excluded). The images were then “skeletonized” and Fiji plug-in “analyse skeleton” was used to obtain information on t-tubule structures as described above.

***Serial Block Face Scanning Electron Microscopy preparation and analysis –***Samples of sheep right atrial appendage (N=3 control, N=3 recovery animals) were fixed in 2.5% glutaraldehyde and 2% paraformaldehyde in 100mM sodium cacodylate buffer, pH 7.2. After washing in sodium cacodylate buffer, samples were prepared as described previously with small modifications (8). Briefly, samples were subsequently stained in: 2% osmium tetroxide and 1.5% potassium ferrocyanide; 1% thiocarbohydrazide; 2% osmium tetroxide, 1% uranyl acetate, and Walton’s lead nitrate with washing in water after each staining step. After staining, samples were dehydrated in an ethanol ascending series (50%, 70%, 90%, 100%, 100%) followed by further propylene oxide dehydration. Increasing concentrations of TAAB 812 hard resin (25%, 50%, 75%, 100%) mixed with propylene oxide were used for infiltration. Finally, samples were embedded in pure resin and cured at 60°C for 36 hours. Samples were extracted from the plastic blocks, glued onto cryo pins, sputter-coated in a gold-palladium alloy and imaged with a Quanta 250 FEG scanning electron microscope, equipped with the Gatan 3View device for serial images collection, and operated at 3.8kV and ~0.46 Torr. Sections were cut at 50 nm and images collected at a nominal resolution of 6.5 -13 nm/px in Gatan .dm4 format. Stacks of images were visualised in Fiji (9) or 3dmod (IMOD) (10) while image segmentation and volumetric analysis was performed in 3dmod.

***Measurement of intracellular calcium -*** Atrial myocytes were loaded with 4µM Fluo-8 AM (Molecular Probes) for 30 minutes. Cells were voltage-clamped using the perforated patch clamp technique AxoClamp2B (Axon Instruments) and pCLAMP software (Molecular Devices, UK) with amphotericin-B (240 μg/ml) in the pipette solution. Cells were stimulated at 0.5Hz using a voltage protocol ramping from -60 to – 40mV and then stepping to +10 mV. Atrial myocytes were imaged confocally using high-speed xyt confocal imaging (Zeiss 7Live; 488 nm excitation and >515 nm emission). Following acquisition of the time series, 20 µM wheat germ agglutinin (WGA) was applied to the cells to visualize the t-tubule and surface membranes and confocal z-stacks were recorded at 210 nm xyz pixel dimensions. Experiments were performed at 37°C.

Changes in [Ca^2+^]_i_ were measured by Script written in Matlab (Mathworks, UK). ROIs, encompassing the cell width, were selected on the xyt confocal time series images. Fluorescence intensity over time for each ROI was plotted and a correction algorithm was applied to the time series to correct for baseline drift between transients. The time series were spilt into individual transients of 400 frames each, background fluorescence was subtracted and fluorescence was normalised (F/F_0_) for each transient. The ‘Dyssynchrony’ analysis algorithm was then used to process the individual transients, where for each pixel of the cell selected, 50% peak fluorescence or half rise time (TF50) and dys-synchrony (standard deviation of the TF50 values) of the systolic rise of [Ca^2+^]_i_ was calculated. Data from three central Ca^2+^ transients in the time series were averaged and used for analysis. To determine the relationship between tubules and Ca^2+^ rise time, the XY coordinates (ImageJ, NIH) for paired data points, distance maps and TF50, were plotted against each other.

***Protein Isolation and Immunoblotting –*** Following removal of the heart (as described above), sheep right atrial appendage tissue samples (N= 7 control, N= 7 HF, N= 7 recovery, animals) were snap frozen and stored in liquid nitrogen until use. Samples were homogenised in RIPA buffer containing protease and phosphates inhibitors (0.1 mg/ml phenylmethanesulphonylfluoride; PMSF, 100 mM sodium orthovanadate, 1 mg/ml aprotonin and 1 mg/ml leupetin). Atrial samples were prepared for SDS-PAGE as described previously (3, 5). Following separation, samples were transferred onto nitrocellulose membranes (GE Healthcare, UK) and blocked using 5% blotto or Superblock (Thermo Scientific, UK). Membranes were incubated 1:1000 with primary antibodies for BIN1, JPH2 (Santa Cruz Biotechnology), MTM1 and Tcap (Abcam). HRP conjugated secondary antibodies (1:20,000, Santa Cruz Biotechnology) were used with chemiluminescent. Protein levels were normalised to an internal standard (IC) which was loaded on all blots. Each sample was repeated in triplicate and data averaged. Additionally, visualisation of Ponceau-S stained membranes ensured even gel loading and transfer. Full length western blots are shown in Supplementary Figure 4.

***Immunocytochemistry–*** Freshly isolated sheep myocytes from the left atrial appendage were plated onto pre-coated μ-slides or laminin coated cell culture slides (Ibidi, GmbH or Fisher Scientific) and left to settle for a minimum of one hour. Cells were fixed in 4% PFA or 100% acetone for 7-10 minutes prior to incubation with WGA Alexa Fluor® 647 conjugate (1:50, Thermo Fisher Scientific, UK) for 2 hours at 4^°^C. Following incubation with WGA, PFA fixed cells were permeabilised with 0.25% Triton x-100 (Sigma) for 10 minutes. Samples were blocked with 10% goat serum followed by incubation with antibodies for NCX (Swant, Switzerland) and RyR (Abcam) (1:100, diluted in 1% serum) overnight at 4°C. Cells were then stained with goat anti-mouse secondary antibodies (A11001 Molecular Probes, 1:500, diluted in 1% serum) conjugated to Alexa Fluor® 488. Following secondary incubation, slides were imaged using a LeicaSP2 confocal microscope (excitation, 488 or 647 nm; emission 500-530 or 663-738 nm). Immunocytochemistry images were digitally deconvolved and thresholded, as described above. Colocalisation analysis was performed (Huygens Professional) to assess the fraction of overlap between NCX, RyR and t-tubules (WGA-Alexa Fluor®) using Manders overlap coefficients.

***Statistics –*** Following enrolment no animals were excluded from the study. Data represent mean ± standard error of the mean (SEM), n cells from N animals. Where data was not normally distributed, data was transformed (log10, reciprocal or square) using a method appropriate to the skew of data (11). Data have been compared using linear mixed model analysis to account for multiple cells from the same animal (SPSS Statistics; IBM, USA). Where comparisons were made on cells, differences were assessed using Student t test or ANOVA (SigmaPlot, Systat Software, USA). GraphPad Prism (GraphPad Software, USA) was used to test for data correlations. Data were considered significant when p<0.05.

### Results

*Echocardiographic parameters in sheep following tachypacing induced heart failure and cessation of pacing.*

Trans-thoracic echocardiograms recorded prior to pacing, at the point of HF and following recovery from HF were used to generate the data in Table 1. Short-axis and long-axis echocardiogram images were used to calculate left ventricular fractional area change; fractional shortening, end diastolic internal diameter (EDID) and end systolic internal diameter (ESID). Fractional shortening = (EDID – ESID) / EDID. Parasternal images from at least three cardiac cycles were averaged for each measurement.

|  | ***Pre-pacing*** | ***Heart failure*** | ***Recovery*** |
| --- | --- | --- | --- |
| **Days paced / recovered** | - | 39.4 ± 3.3 | 37.3 ± 1.2 |
| **End Diastolic Internal Dimension (cm)** | 2.88 ± 0.15 | 4.18 ± 0.22 *** | 3.56 ± 0.20^#,^ * |
| **End Systolic Internal Dimension (cm)** | 1.24 ± 0.11 | 3.32 ± 0.25 *** | 2.11 ± 0.19 ^##,^ * |
| **Fractional area change (short-axis view)** | 0.69 ± 0.01 | 0.30 ± 0.04 *** | 0.58 ± 0.03 ^###,^ * |
| **Fractional shortening (long axis view, m-mode imaging)** | 0.58 ± 0.03 | 0.21 ± 0.03 *** | 0.39 ± 0.03 ^###,^ *** |
| **Cell width (µm)** | 16.3 ± 0.64 | 18.62 ± 0.20 * | 15.8 ± 1.07^#^ |

**Supplementary Data, Table 1. Cardiac function was improved following cessation of rapid pacing in sheep.** Mean data summarising echocardiographic parameters in sheep. * *P*<0.05, *** *P*<0.001 vs Control; ^#^ *P*<0.05, ^##^ *P*<0.01, ^###^ *P*<0.001 vs HF, using One-way ANOVA with repeated measures. Measurements were taken at each time point for *n* = 11 animals (8 paired).

*Atrial t-tubules were characterised based on morphology*

The detailed three-dimensional structure of atrial t-tubules was determined using serial block face Scanning Electron Microscopy (sbfSEM). Both control and recovered atrial t-tubules adopted distinct morphologies which were categorized into 12 groups. The criteria for these groups are given in Table 2 and typical examples are shown in Supplemental Figure 1A (t-tubule images orientated such that they connect with the cell membrane at the base of the image).

| *T-tubule morphology* | *Description* |
| --- | --- |
| Column | Uniform in width and long in length. |
| Angled | Uniform in width and long in length but bent at top (90° ± 45). |
| Stump | Short in length. |
| Club | Narrow at base, at least twice as wide at head. |
| Intermediate | Mid way between stump and fully developed t-tubule. |
| Cactus | Column with 1 branch - branch is shorter than and usually at an angle to main trunk. |
| Branched | More than 1 branch and the branch can go in any direction. |
| Pair | Either the t-tubule splits to run either side of z line or t-tubule is associated with another structure on same z line. |
| Random | Other – when t-tubule does not fit into the other categories. |
| Oak tree | Extremely branched with longitudinal elements. |
| Longitudinal | Runs across several sarcomeres with no main transverse trunk. |
| Lattice | Grid like pattern. |

**Supplementary Data, Table 2. T-tubule morphology criteria.** Reconstructed tubule morphology was defined according to the criteria in Table 2.

*T-tubule morphology was altered following recovery from heart failure in atrial myocytes.*

SbfSEM revealed control atrial t-tubules are not uniform in structure but adopt a range of morphologies (Supplemental Figure 1Ai), the most common of which are ‘column’ and ‘angled’ t-tubules (Supplemental Figure 1B). Following recovery from HF, fewer t-tubules adopt these normal morphologies with many being exceptionally disordered displaying extreme morphologies such as ‘oak tree’, ‘random’, ‘longitudinal’ and ‘lattice’ (Supplemental Figure 1Aii & B). These extreme morphologies were largely restricted to recovery cells (Supplemental Figure 1Bi&ii) constituting 61% of the total t-tubule volume in these cells. T-tubule doublets also occurred more frequently in recovered cells, possibly explaining why, compared with control, more recovered t-tubules arise in locations other than the z-line and are often seen to span the sarcomere (Supplemental Figure 1 Ci&ii).

Our data also suggests that following recovery from HF an association exists between disordered mitochondria and t-tubule structure. Here mitochondria form disordered, wide beds and often appear out of place e.g. they can occur on the z-line as shown in Supplemental Figure 2 where the recovered t-tubule branches to pass either side of the mitochondrion.

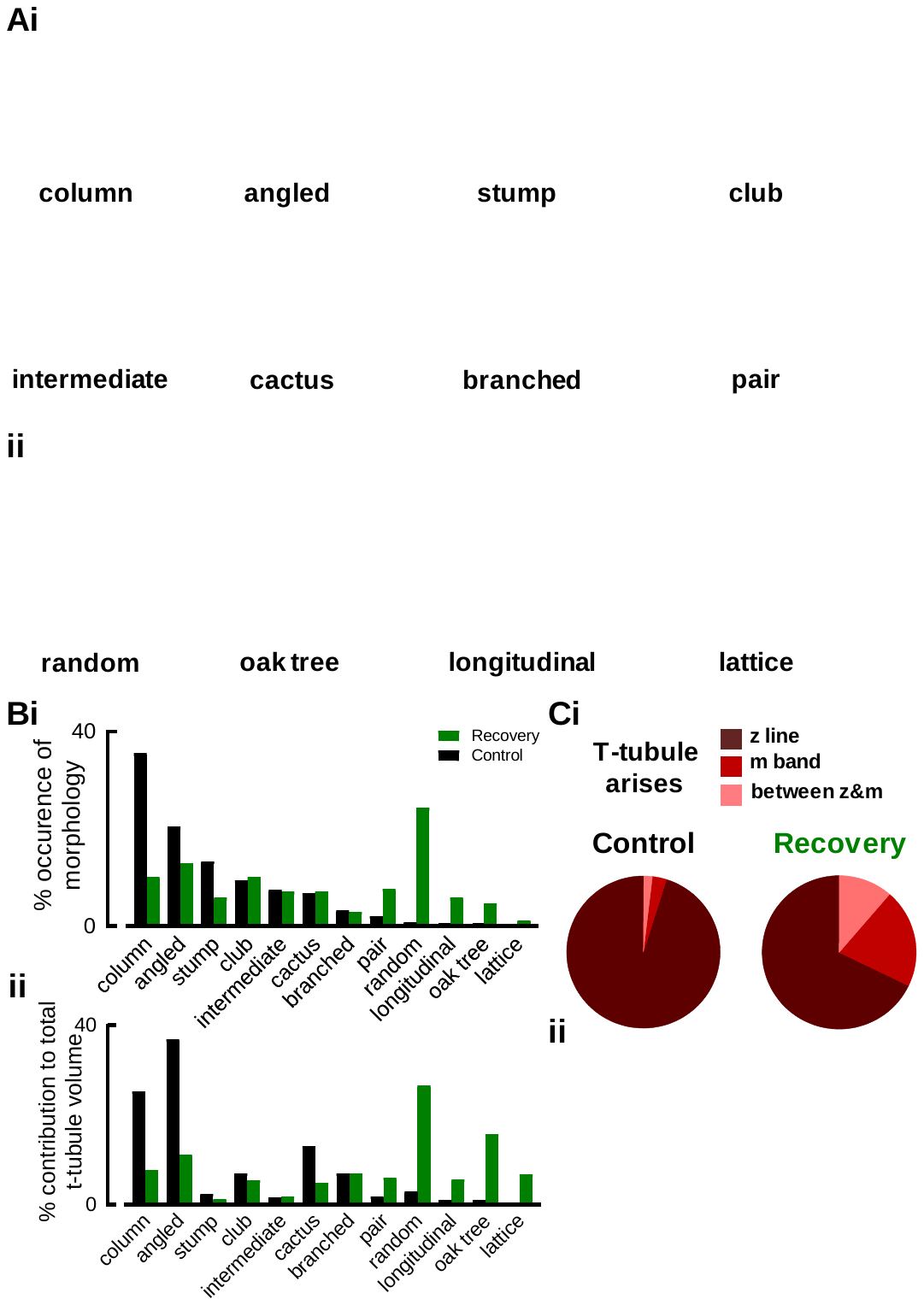

**Supplementary Data, Figure 1. T-tubule ultrastructure following recovery from heart failure. (A)**3D reconstructed representative images of the different t-tubule morphologies described in Table 2, scale bar denotes 2µm. (Ai) represents morphologies common in the control atria whereas (Aii) depicts disordered t-tubules common in recovered atrial myocardium. (B) Summary data showing the % occurrence (i) and % contribution to the total t-tubule volume (ii) of each t-tubule morphology in control (black) and recovered (green) atrial myocytes, n= 7-9 cells from N=3 animals per group. (C) T-tubules arise at non z-line sites more often following recovery from HF. (Ci) Summary data showing the sites where t-tubules arise. (Cii) T-tubules can span sarcomeres; 3D reconstruction from a sbfSEM image, z-lines are shown in grey and t-tubules shown in red, scale bar denotes 5µm.

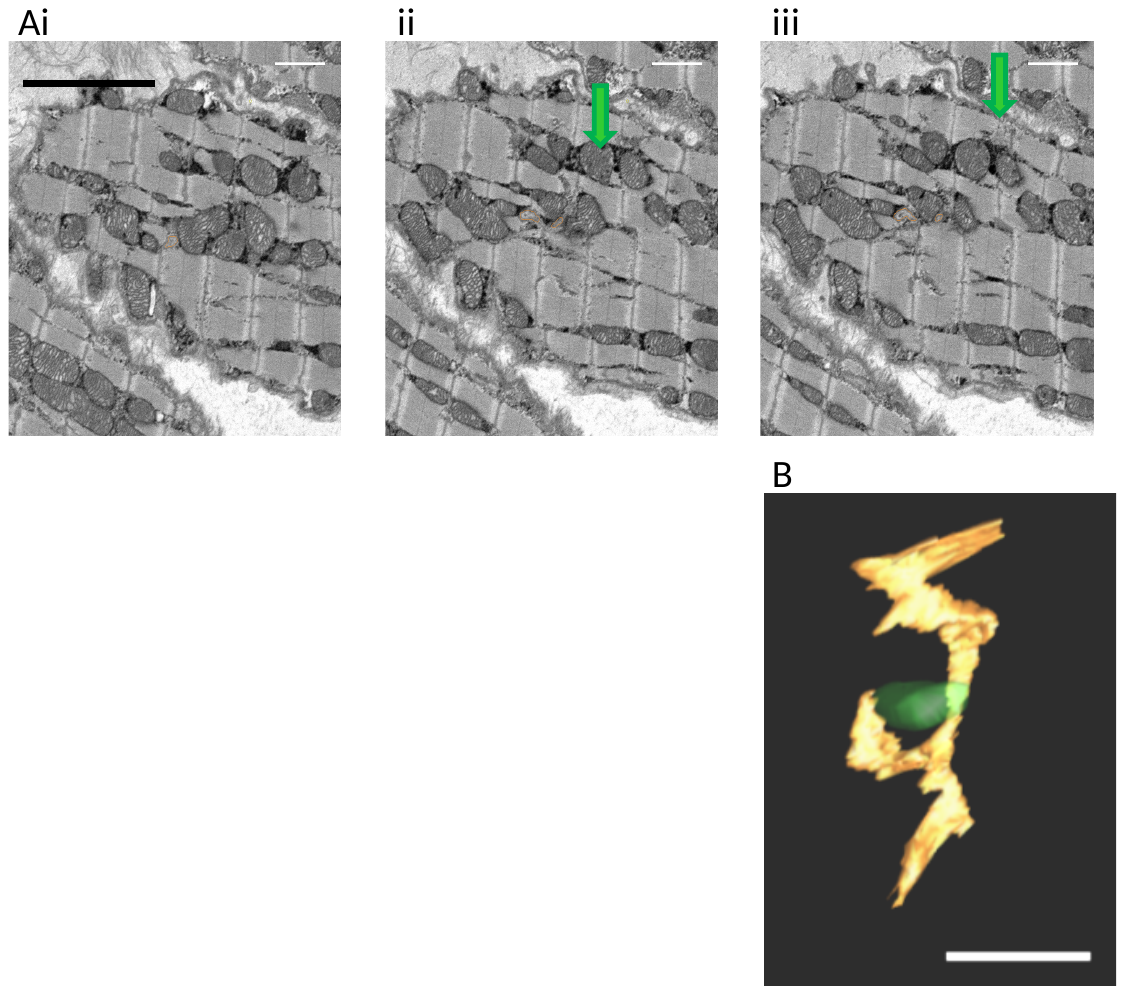

**Supplementary Data, Figure 2. Mitochondrial disorder correlates with t-tubule structure following recovery from HF.**(A) Image sequence to show a recovered atrial t-tubule (orange) move along the Z-line (i) and branch around a mitochondrion situated on the z-line (green arrows).(B) Reconstruction of the orange t-tubule and green mitochondrion from (A) to show the t-tubule branching around the mitochondrion; scale bars are 2µm.

*Cellular distribution of calcium handling proteins*

Immunocytochemistry was used to determine if dyadic structure was altered in HF and following recovery from HF. The Na^+^ / Ca^2+^ exchanger (NCX) colocalised with the t-tubules and the surface sarcolemma membrane in the control and recovery cells (Manders coefficients,Huygens Professional). In the HF cells, where t-tubules were almost completely absent, the NCX staining was only present on the cell surface sarcolemma (Supplemental Figure 3Ai-ii).

RyRs occurred on the z-line in control cells but were sparse at the surface (Supplemental Figure 3Bi and Biv). RyRs localized with the cell membrane at the t-tubules (WGA, red; supplemental Figure 3Bi, total merge, t-tubule merge & Bii-iii). Colocalisation between t-tubules and RyRs was not absolute in control cells as atrial t-tubules do not occur at every z-line. In HF, t-tubules were absent, RyR staining fragmented (Supplemental Figure 3Bi) and colocalization between RyRs and cell membrane decreased (Supplemental Figure 3Bi t-tubule merge and Bii-iii). However, RyR colocalization with the surface membrane increased (surface merge and Biv) suggesting there are more RyRs at the cell surface in HF. Following recovery from HF RyRs adopted a z-line localization being sparse on the surface membrane, as in control cells (Supplemental Figure 3Bi). In recovery cells colocalization between RyRs and cell membrane returned to control levels since colocalization occurred where disordered t-tubules bisect the z-line (Supplemental Figure 3Bi-iv).

Compared to control and recovery, where confocal imaging shows continuous RyR staining, in HF RyR staining was fragmented and occurred between z-lines (Supplemental Figure 3 Ci). RyR distribution was measured firstly, using the Fiji algorithm “analyse skeleton” to determine how many separate areas of RyR staining existed along the z-line. The summary data shows that the average number of RyR stained regions per sarcomere increased in HF and returned to control values following recovery (Supplemental Figure 3Cii). Secondly, RyR intensity plots, across RyR stained sarcomeres displayed disordered patterns across both the longitudinal (blue dashed line on Ci) and transverse directions (orange dashed line on Ci) in HF (Supplemental Figure 3 Ciii). This data suggests that as well as fragmentation along the z-line RyRs occur between z-lines in HF atrial cells. Recovery from HF led to the general distribution of the RyRs to returning to control levels.

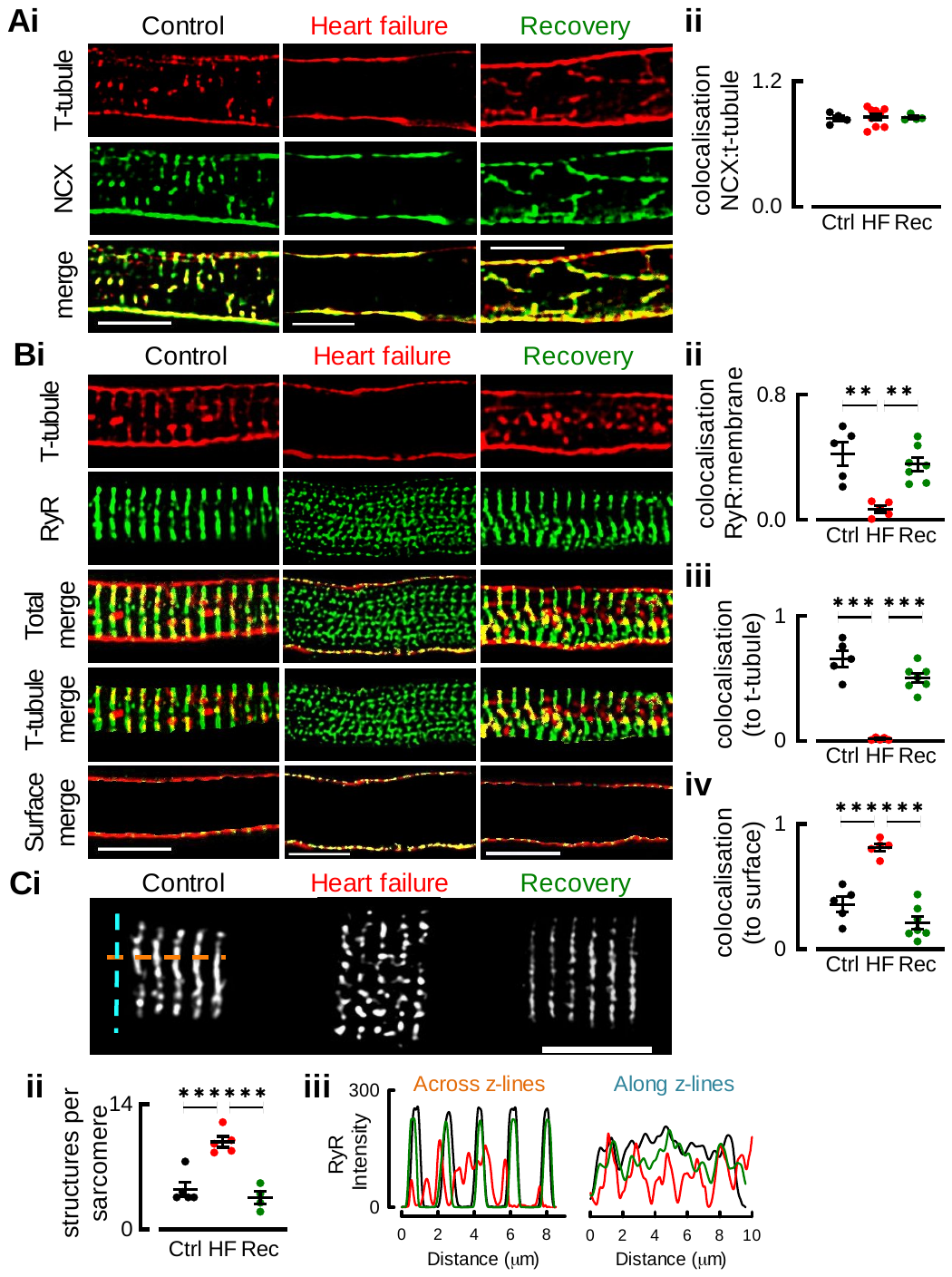
**Supplementary Data, Figure 3. Cellular distribution of Ca handling proteins in control, heart failure and recovery atrial myocytes**.(Ai) Immuno staining of NCX (green) and t-tubule or surface membrane (red) in atrial myocytes, yellow represents colocalisation. (Aii) Mean data showing Manders colocalisation of NCX and t-tubules in control, HF and recovery myocytes. (B) Immuno staining of RyR (green) and t-tubules (red), yellow represents colocalisation (i) and mean data summarizing Manders colocalisation of RyR and cell membranes (ii), RyR and t-tubules (iii) and RyR with surface membrane (iv). (C) RyR distribution along the sarcomere (i) and mean data for number of RyR structures per sarcomere (ii) and representative intensity blots (iii) across RyR stained sarcomeres from (i) in both longitudinal and transverse directions. N= 2 animals per group (NCX, n= 4-9 cells; RyR, n= 5-7 cells), ** p<0.01, *** p<0.001 vs ctrl, ## p<0.01, ### p<0.001 vs HF tested by ANOVA, scale bars =10µm.

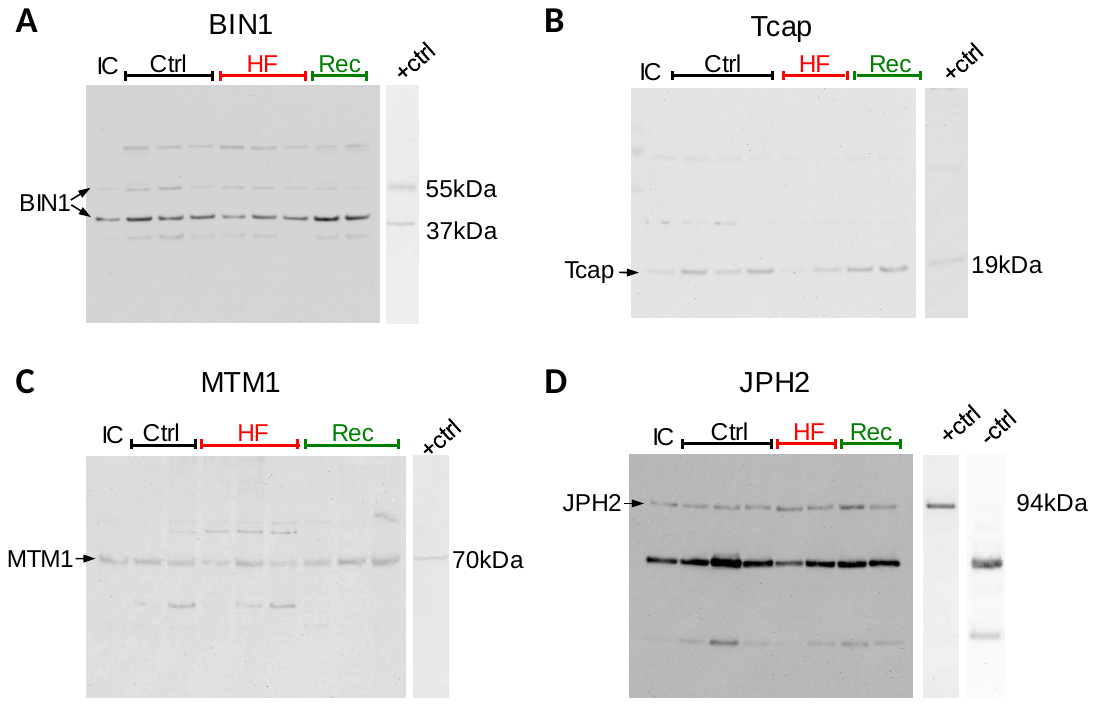

**Supplementary Data, Figure 4. Full length Western blots for BIN1, Tcap, MTM1 and JPH2.** Controls were as per the antibody data sheet and the negative control for JPH2 is the secondary antibody only.
